# R Functions for Analysis of Continuous Glucose Monitor Data

**DOI:** 10.1101/625137

**Authors:** Tim Vigers, Christine L. Chan, Janet Snell-Bergeon, Petter Bjornstad, Philip S. Zeitler, Gregory Forlenza, Laura Pyle

**Author notes:** Corresponding Author (LP).

## Abstract

Continuous glucose monitoring (CGM) is an essential part of diabetes care. Real-time CGM data are beneficial to patients for daily glucose management, and aggregate summary statistics of CGM measures are valuable to direct insulin dosing and as a tool for researchers in clinical trials. Yet, the various commercial systems still report CGM data in disparate, non-standard ways. Accordingly, there is a need for a standardized, free, open-source approach to CGM data management and analysis. Functions were developed in the free programming language R to provide a rapid, easy, and consistent methodology for CGM data management and analysis. Summary variables calculated by our package compare well to those generated by various CGM software, and our functions provide a more comprehensive list of summary measures available to clinicians and researchers. Consistent handling of CGM data using our R package may facilitate collaboration between research groups and contribute to a better understanding of free-living glucose patterns.

## Introduction

Continuous glucose monitoring (CGM) technology has transformed diabetes care over the past 15 years by allowing clinicians to measure free-living glucose patterns. During this period, CGM use has increased from < 5% of patients to almost 50% in some age groups [1]. With recent reports detailing the benefits of CGM time in range metrics as predictive of long-term vascular outcomes [2] and as an indicator of glucose management or estimated hemoglobin A1c (HbA1c) [3], CGM use will likely continue to increase in both research and clinical settings. Despite the increasing use of CGM for treatment and research, a standardized, free, open-source approach to data management and analysis is lacking [4].

CGM manufacturers use proprietary algorithms to create reports and calculate summary measures for patients and clinicians. As a result, it may be difficult to compare results obtained using different CGM devices and to understand the sources of variability that could influence CGM outcomes. In addition, research questions may require summary measures that are not available in accompanying reports (e.g., use of a different cut-point for hyperglycemia). Furthermore, use of the summary values provided by each CGM platform sometimes requires that data be entered by hand into a database or spreadsheet prior to analysis. This is a time-consuming and error prone process that will benefit from automation. The use of a free and open source program to analyze raw sensor glucose values will enable researchers to define their own variables of interest and standardize calculation of summary measures across different CGM devices.

There have already been a few attempts to develop such systems, including the EasyGV macro-enabled Excel workbook [5], AGP Report (agpreport.org), and Tidepool (tidepool.org). However, there are reports suggesting that EasyGV poorly matches other calculations of mean amplitude of glycemic excursion (MAGE) [6], and it does not permit the various definitions of a significant excursion (i.e. greater than 1 standard deviation (SD), 2 SDs, etc.). Although Tidepool appears to be an excellent option for patients and clinicians, it is not free for use in research, and many smaller investigator-initiated studies cannot afford the additional expense. Also, their open source code requires significant coding knowledge in multiple programming languages which limits accessibility and widespread use.

To address this need, we have developed a package written entirely in the statistical programming language R (R Foundation for Statistical Computing, Vienna, Austria). The package currently works with data from Diasend (www.diasend.com), Dexcom (www.dexcom.com), iPro 2 (http://professional.medtronicdiabetes.com/ipro2-professional-cgm), Libre (www.freestylelibre.us), and Carelink (www.medtronicdiabetes.com/products/carelink-personal-diabetes-software), with plans to add support for other platforms as CGM technology advances. Additionally, data can be manually formatted to work with these functions if necessary. The package is available on The Comprehensive R Archive Network (CRAN) under the name ‘cgmanalysis’ and the source code can be found at https://github.com/childhealthbiostatscore/R-Packages, which allows for version control and forking if users need to alter functionality, and includes a short user guide for those with limited R experience.

### Summary Measures of Glycemia

Although CGM is not a new technology, there is still debate regarding the advantages and disadvantages of various CGM metrics for use in clinical care and as research outcomes. The American Diabetes Association (ADA) recently proposed a set of key metrics for CGM analysis [7], all of which are calculated by our code, in addition to the glucose management index (GMI) [3], time in range [2], and other variables proposed by Hernandez et al.[8]. An easy method to calculate these important summary variables from a variety of sources of CGM data has the potential to contribute to the standardization of the use of these metrics. A list of summary variables produced by our default code is available in Table 1, and Table 2 provides comparisons between the package and proprietary software. The code can be easily modified to include further variables of interest, to be released in future version updates. Further, because the package is open source, individual users can create their own modifications.

**Table 1:**
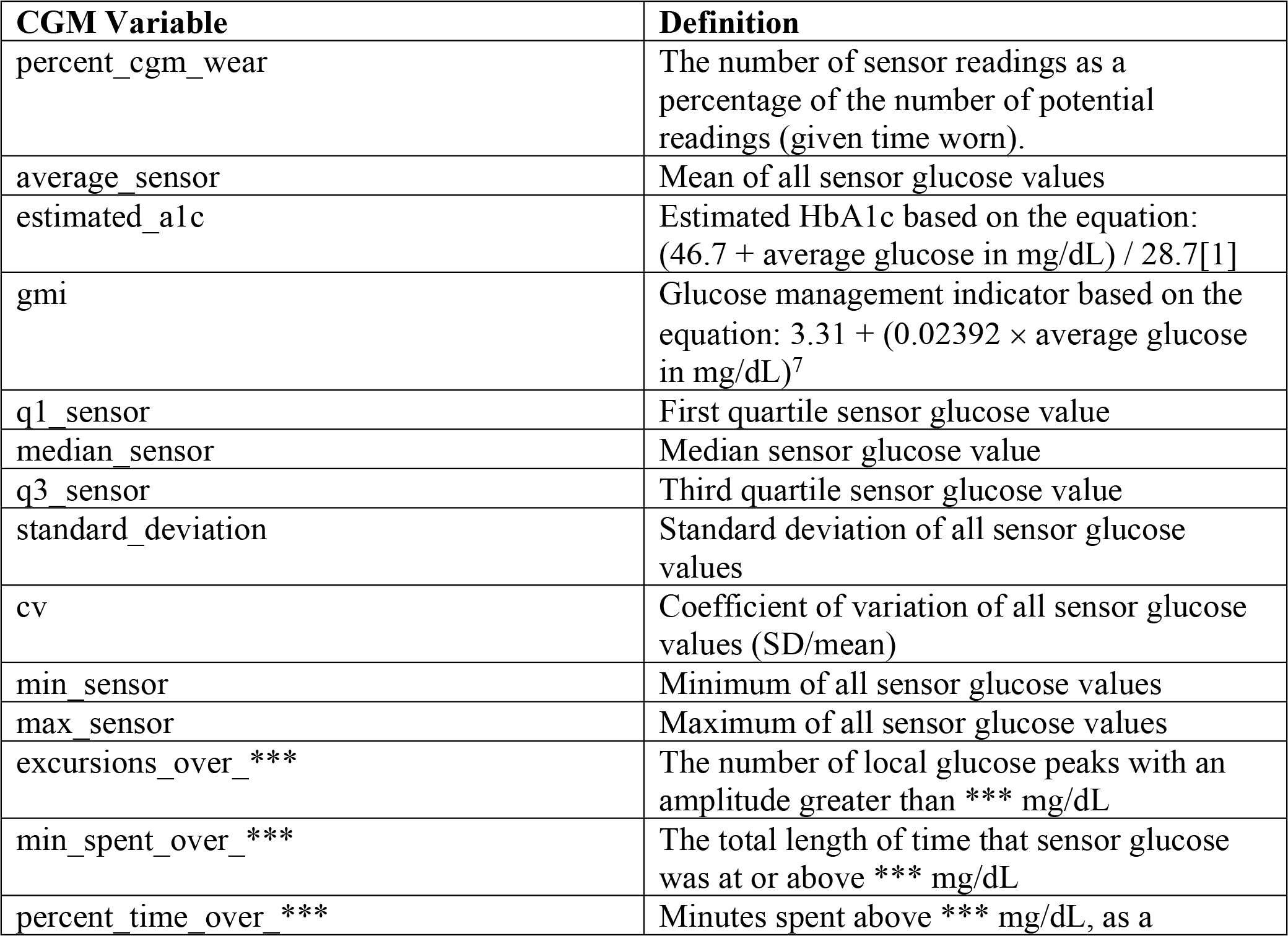
Summary Measures of Glycemia.

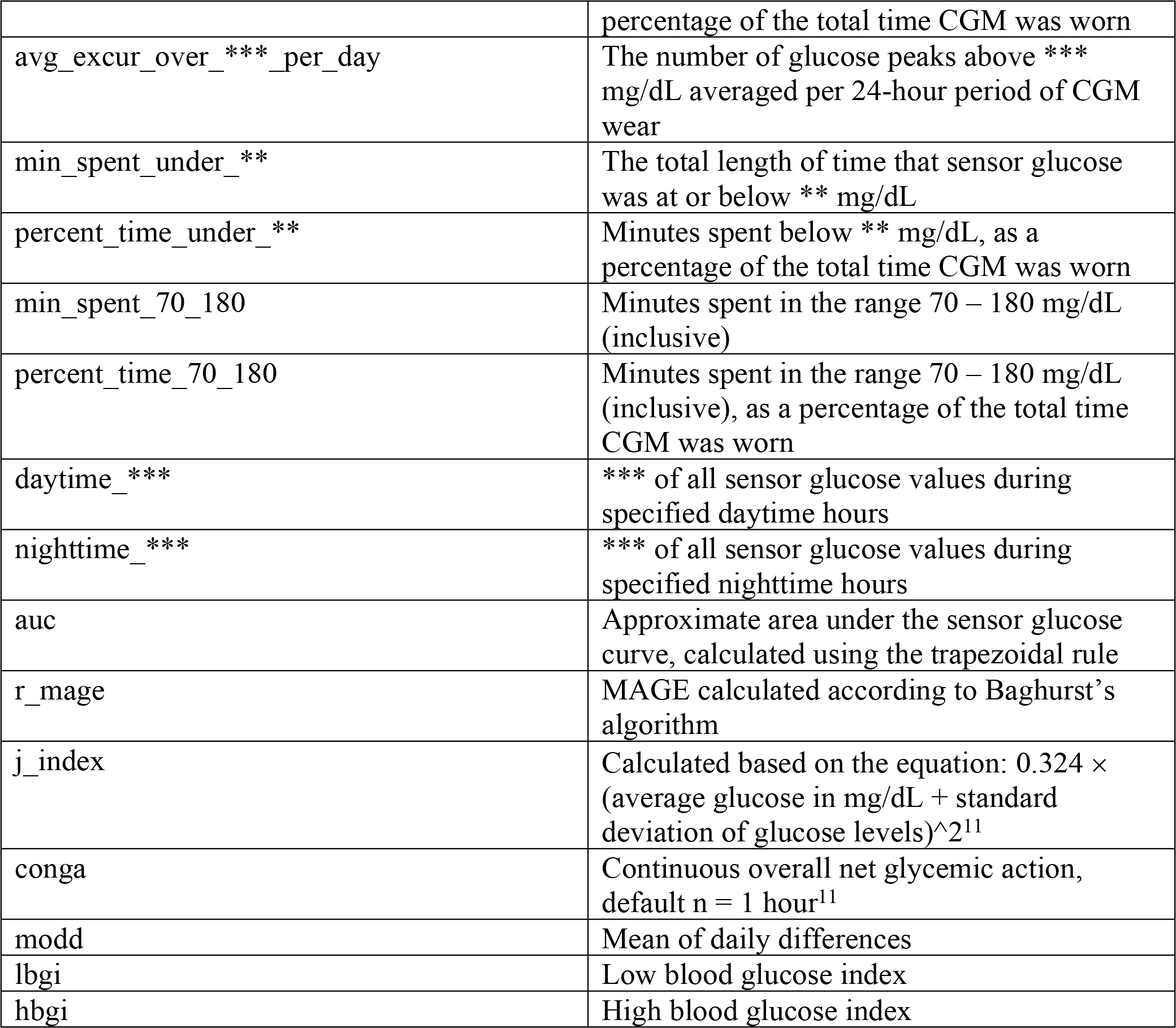

**Table 2:** Summary Variable Comparisons.

A. iPro 2 software (high excursion defined as > 140 mg/dL for 15 minutes, low defined as < 60 mg/dL for 15 minutes)
|  | cgmanalysis | iPro |
| --- | --- | --- |
| # Sensor Values | 2000 | 2000 |
| Highest | 282 | 282 |
| Lowest | 70 | 70 |
| Average | 126.87 | 127 |
| Standard Dev | 30.79 | 31 |
| # High Excursions | 31 | 32 |
| # Low Excursions | 0 | 0 |
| % Time Above 140 | 24.85 | 24 |
| % Time Below 60 | 0 | 0 |

B. Carelink 670G
|  | cgmanalysis | Carelink 670G |
| --- | --- | --- |
| Average | 123.65 | 124 |
| Standard Dev | 37.53 | 38 |

C. Dexcom Clarity
|  | cgmanalysis | Dexcom Clarity |
| --- | --- | --- |
| Average | 175.68 | 176 |
| Standard Dev | 67.10 | 68 |
| Time in Range | 55.66 | 56 |

D. Diasend
|  | cgmanalysis | Diasend |
| --- | --- | --- |
| # Sensor Values | 184 | 184 |
| Highest | 411 | 411 |
| Lowest | 54 | 54 |
| Average | 193.23 | 193 |
| Standard Dev | 89.67 | 89 |
| Values above 200 | 44.57% | 44.57% |

## Methods

### Package Design

Our package consists of three simple functions: cleandata(), cgmvariables(), and cgmreport(). The data cleaning function iterates through a directory of CGM data exports and produces new files that then serve as input to the CGM variable calculator and the CGM report generator. The initial directory can contain files from different sources, as the function identifies the relevant timestamp and glucose values for each file format. By default, the cleaning function will fill in gaps in glucose data less than 20 minutes long using linear interpolation. It will also remove 24-hour periods containing gaps larger than 20 minutes, so that there will be an equal number of daytime and nighttime values, important for calculating some variables, such as AUC. The user can specify a different maximum gap to fill by interpolation and can also choose whether to remove days with larger gaps. Ideally, the CGM data should be exported and then cleaned using this package, and not manually edited. However, if a file does require manual data editing, these functions will work on the three-column format detailed in the package documentation.

Once the data have been cleaned, the CGM variables described in Table 1 are calculated using the cgmvariables() function. By default, blood glucose must be above a threshold for at least 35 minutes or below a threshold for at least 10 minutes to count as an excursion, but these parameters can be changed by the user if necessary. Likewise, daytime (e.g. for daytime vs. nighttime AUC or maximum glucose) is defined as 6:00 to 22:00 by default, but these can be set depending on user needs. MAGE is calculated using Baghurst’s algorithm [9], which we have coded in R. By default, the function includes blood glucose excursions greater than 1 SD from the mean in calculation of MAGE, but there are options for 1.5 SD and 2 SD as well.

Our code was originally written to produce data tables for upload to a Research Electronic Data Capture (REDCap) database [10], which influenced the selection of variable names in the final output. These names can be changed in the code itself or by simply editing the function’s output. These variables are stored in separate columns of a new data frame (the function’s output), with each record identified by the patient ID.

In addition to producing calculated variables, our package can also plot CGM data in a few ways. First, the function concatenates all the CGM data in the specified directory into one data table and plots the aggregate data in the style of the standard AGP report (http://www.agpreport.org), the aggregate daily overlay (ADO). This method uses Tukey smoothing after rounding each timepoint to the nearest 10-minute mark, then plots the median, inter-quartile range, and 5 and 95 percentiles at each time of day (with plans to add more options in the future). The package also produces a similar aggregate plot with a Loess-smoothed (locally estimated scatterplot smoothing) average overlaid on points representing every single glucose value. For smaller data sets, this type of plot gives a meaningful overview of daily glucose trends. Finally, the third type of plot uses a Loess-smoothed average for each patient with glucose values color-coded by participant.

### Comparison of cgmanalysis package and proprietary software

Our functions were compared to proprietary CGM software using clinically collected data from iPro 2, Carelink 670G, Dexcom Clarity, and Diasend. The data were exported from each platform, formatted using the cleandata() function, then summarized using the cgmvariables() and cgmreport() functions. The data were not cleaned prior to plotting and summary variable calculation, and summary variable parameters were altered from default (e.g. defining an excursion as 15 minutes above or below threshold for iPro 2 data) in order to better match the CGM results. Because each CGM device provides different and limited summary variables, we were only able to compare a small subset of our package’s output and were not able to directly test more complex variables, such as MAGE or CONGA.

## Results

Fig 1 is an example of the ADO plot made using approximately 25,000 simulated CGM values, and Fig 2 is the version of the ADO with Loess smoothing, using the same data as in Fig 1. Fig 3 is the patient-specific plot, made with a subset of the simulated data.

**Fig 1:**
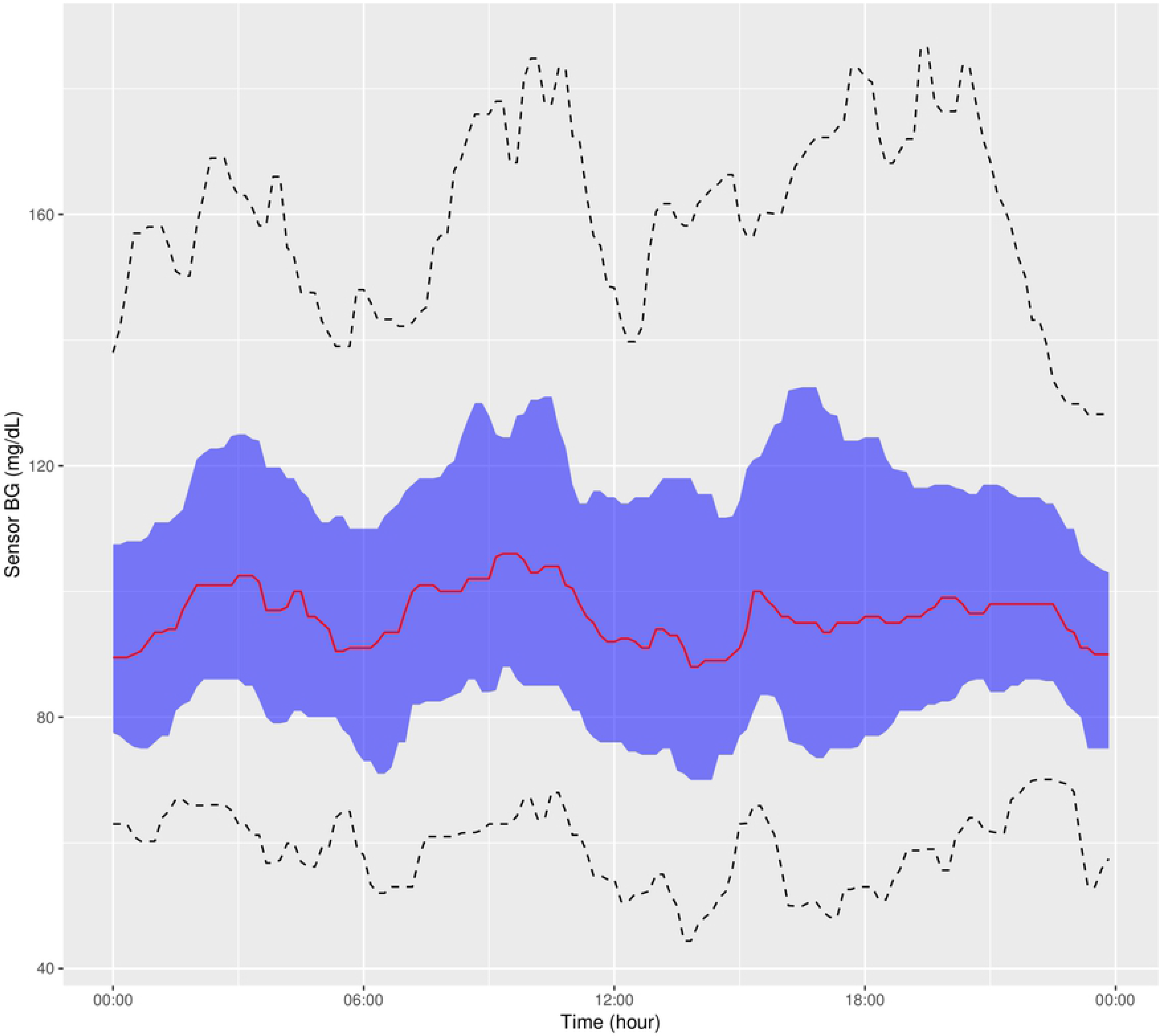
Aggregate Daily Overlay (Tukey Smoothing)

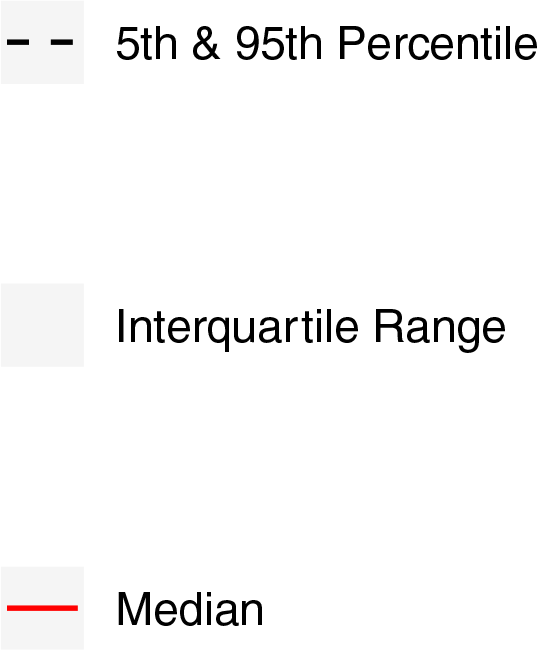

**Fig 2:**
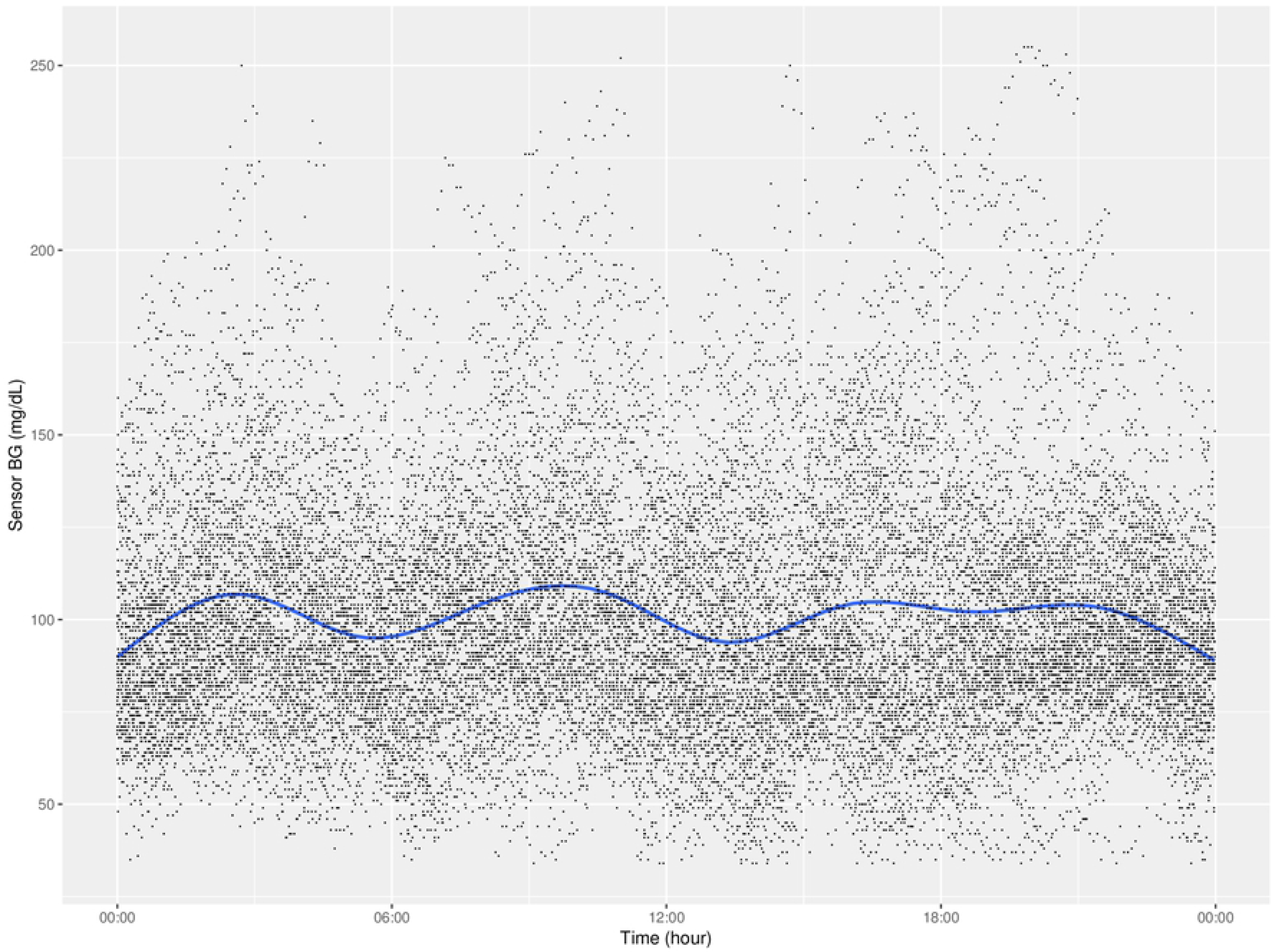
Aggregate Daily Overlay (Loess Smoothing)

**Fig 3:**
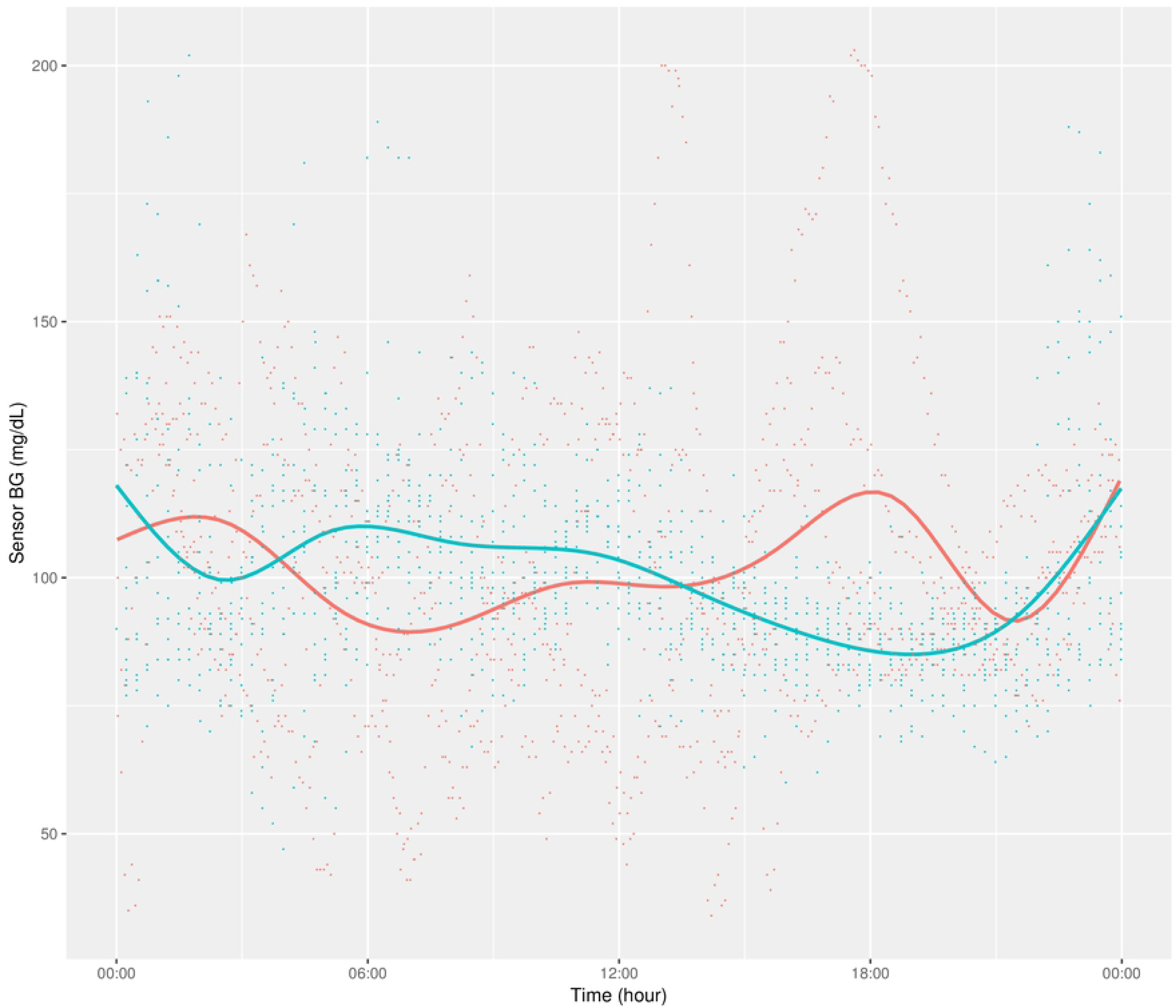
Daily Overlay per Subject (LOESS Smoothing)

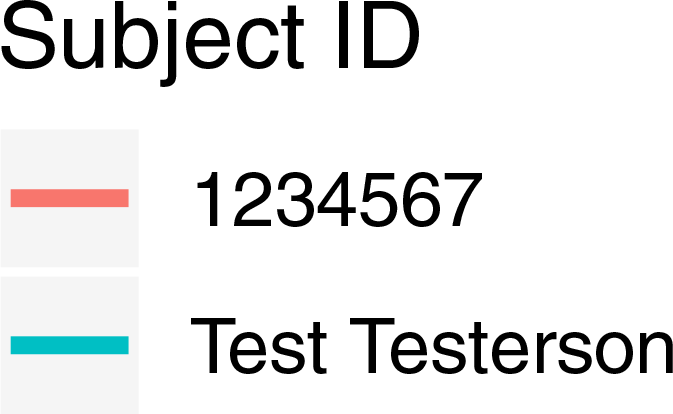

Table 2 shows the results of summary variable comparisons between four different proprietary CGM devices and our cgmanalysis package. Most of the differences in these comparisons are small and the result of rounding. Overall the package appears to be capable of reproducing proprietary calculations when run with non-default settings, although in the comparison to the iPro 2, there was a difference of 1 high excursion.

Figs 4a-d show the comparisons of the graphical outputs produced by the proprietary software and the cgmanalysis package. In the graphs produced by the cgmanalysis package, glycemic patterns at each hour of the day are clearly visible and match the CGM device outputs well. However, some of the proprietary software appear to apply different smoothing algorithms, resulting in slightly different patterns across time.

**Fig 4a:**
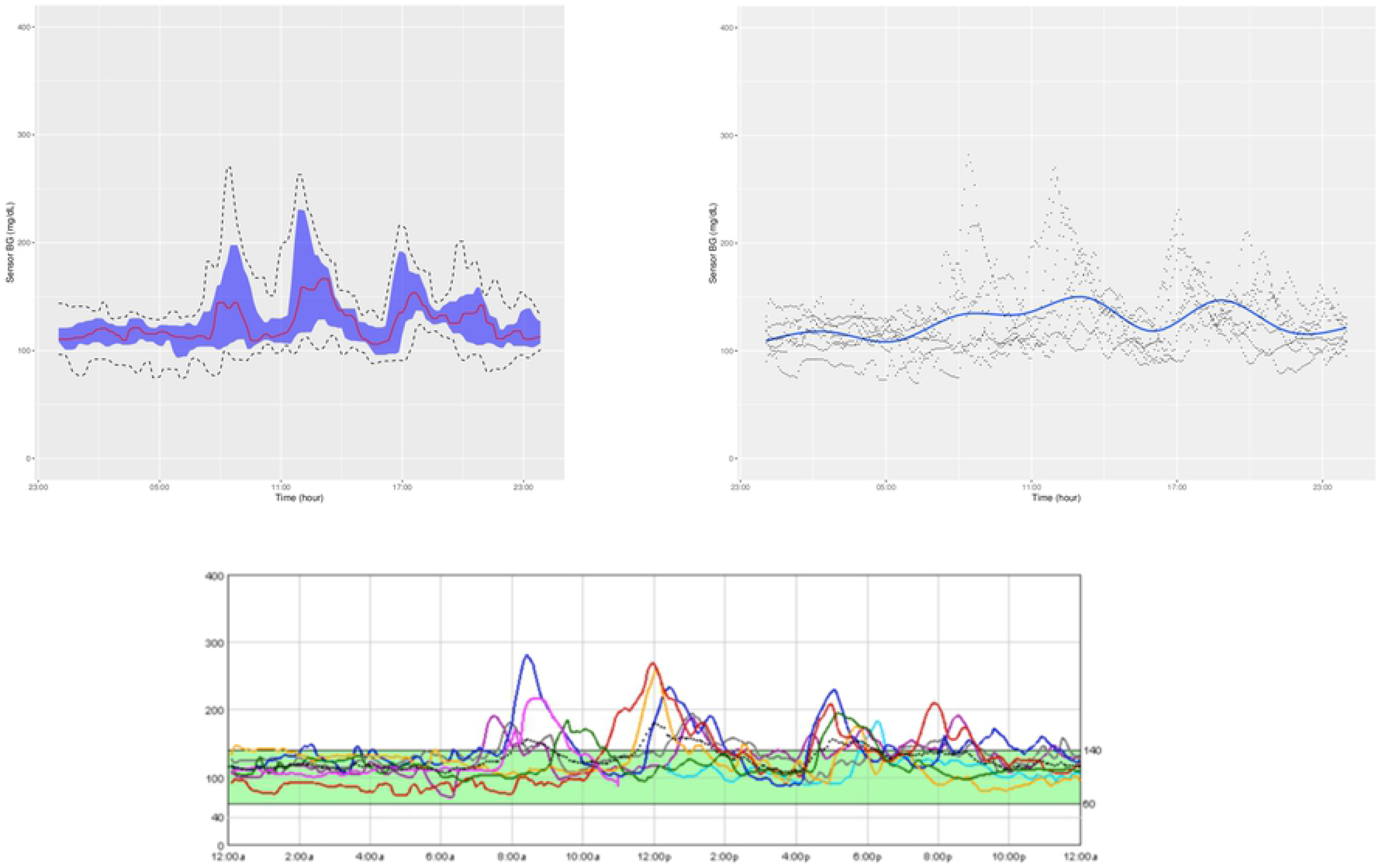
“cgmanalysis” Package Plots Compared to iPro 2 Daily Overlay. Clockwise from top left: Aggregate Daily Overlay (Tukey Smoothing), Aggregate Daily Overlay (Loess Smoothing), iPro 2 Daily Overlay

**Fig 4a.**
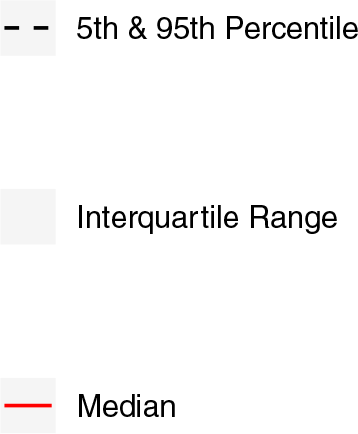
Tukey AGP (Top Left) Legend.

**Fig 4b:**
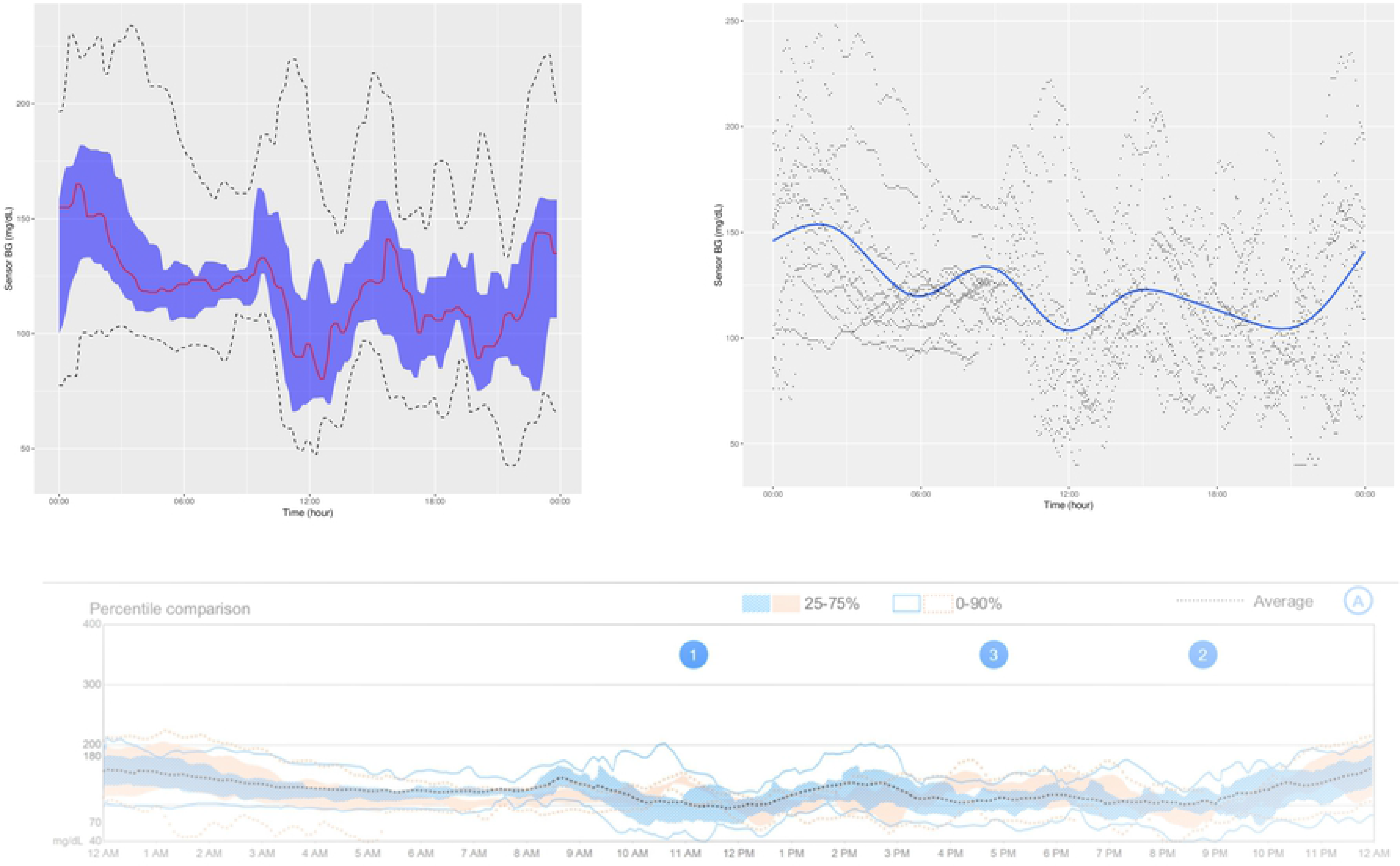
“cgmanalysis” Package Plots Compared to Carelink 670G Daily Overlay. Clockwise from top left: Aggregate Daily Overlay (Tukey Smoothing), Aggregate Daily Overlay (Loess Smoothing), Carelink 670G Daily Overlay

**Fig 4b.**
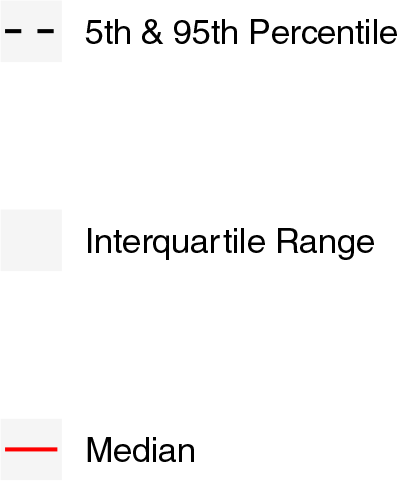
Tukey AGP (Top Left) Legend.

**Fig 4c:**
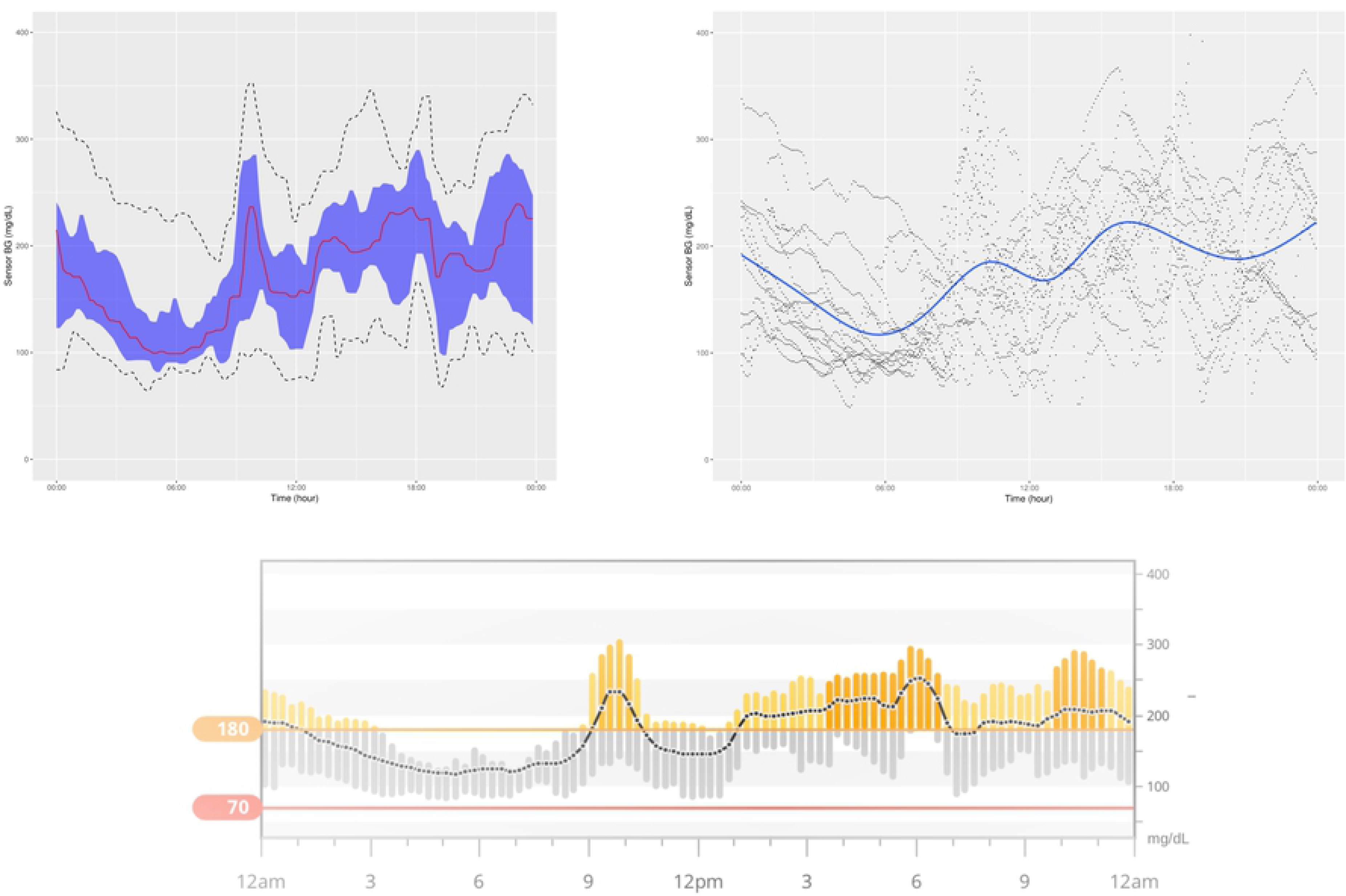
“cgmanalysis” Package Plots Compared to Dexcom Clarity Daily Overlay. Clockwise from top left: Aggregate Daily Overlay (Tukey Smoothing), Aggregate Daily Overlay (Loess Smoothing), Dexcom Daily Overlay

**Fig 4c.**
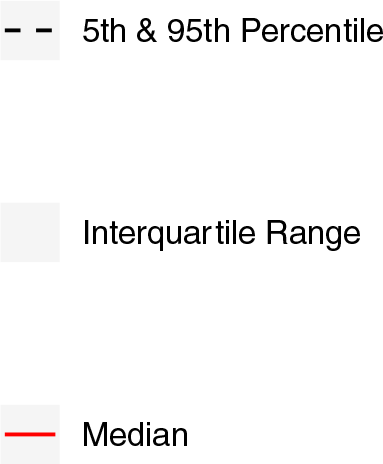
Tukey AGP (Top Left) Legend.

**Fig 4c.**
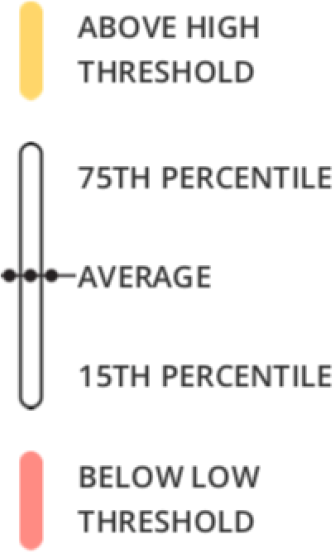
Dexcom Clarity (Bottom) Legend.

**Fig 4d:**
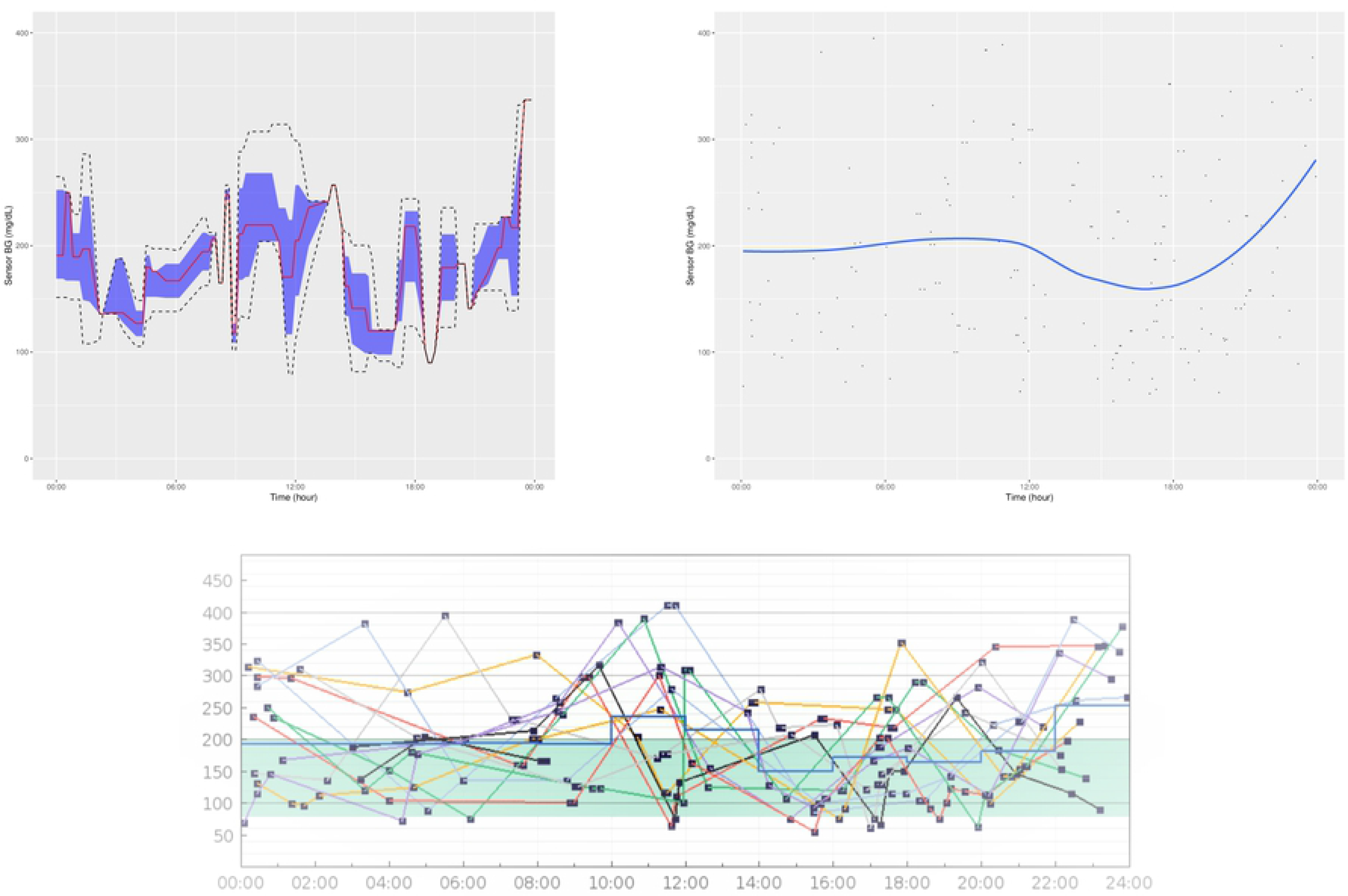
“cgmanalysis” Package Plots Compared to Diasend Daily Overlay. Clockwise from top left: Aggregate Daily Overlay (Tukey Smoothing), Aggregate Daily Overlay (Loess Smoothing), Diasend Daily Overlay

**Fig 4d.**
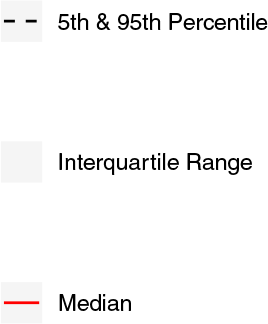
Tukey AGP (Top Left) Legend.

**Fig 4d.**
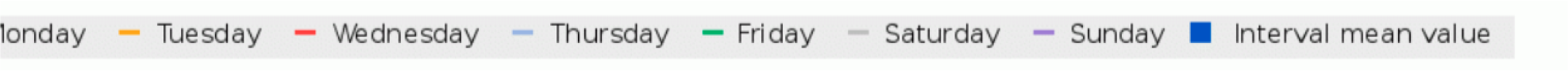
Tukey AGP (Bottom) Legend.

## Discussion

The summary variables produced by the cgmanalysis package match those from the proprietary software for all platforms assessed, and differences are mainly due to rounding discrepancies. Compared to the iPro 2, the number of high excursions differed by 1. Without access to the iPro algorithms we are unable to determine why these counts disagree, but the difference is not likely of clinical significance. The graphical outputs from the cgmanalysis package are similar to the CGM device output in terms of the glycemic patterns by hour of day, although there are small differences, likely due different smoothing algorithms.

There are several limitations to our comparison of the cgmanalysis package to the proprietary software output. CGM devices only calculate a few summary variables, and accordingly it is difficult to test this package cohesively. Also, gold standard calculations do not exist for many of these variables, which makes verifying our results difficult. We hope that by making this package freely available and open source, these limitations will be minimized through widespread testing. Perhaps the greatest limitation to the software itself is the lack of an easy to use graphical user interface (GUI), which may prevent its use by clinicians with limited programming experience. We have included detailed documentation in the CRAN package, as well as a new-user guide on GitHub, but using the package still requires enough technical knowledge that it may be inaccessible to some users. None of the authors are software engineers, and the package is undoubtedly less efficient than it could be. Again, we hope that the free and open source nature will contribute significantly to improving the code over time, both as a result of outside contributions and our own planned updates.

In conclusion, our software provides a standardized, free, open-source approach to manage and analyze CGM data, enabling sharing of data across technology platforms, collaboration between research groups, and more effective use of the growing pool of CGM data. The advantage of using R functions rather than licensed statistical software, or a web-based or desktop application, is that R is freely available and open source. Clinicians or investigators can alter the code according to their needs and anyone can contribute to the development of the program, as CGM research and technology advance.

